# Remote sensing tools for plant invasions assessment in native tropical forests of a volcanic island

**DOI:** 10.1101/2022.05.15.491984

**Authors:** Roussel Guillaume, Flores Olivier

## Abstract

Surveying and monitoring biological invasions by plants can greatly benefit from the application of remote sensing techniques. In tropical insular enviroments however, multiple challenges makes the task difficult. Invasive species can be diverse and many, so that evaluating an overall invasion level may help to distinguish uninvaded native forests from forests invaded at various levels. Also, accessibility can limit the availability of field-based data documenting the ground truth of plant invasions. Here we present a methodological study with the doublefold objective of first estimating an overall index of invasion and level of native forest canopies and second detect two particular exotic (*Cinamomum camphora*, Lauraceae and *Cryptomeria japonica*, Taxodiaceae) planted in monospecific stands. The index of overall invasion level comprises four classes derived elsewhere from field observations and expert knowledge. A thematic map provided polygons representing monospecif plantations. Textural features of sub-regional regions of interest and polygons were derived using Haralick coefficients on co-occurrence matrices, and morphological profiles. Classification algorithms were applied to produce invasion classes (k-means) or detect target species (Support Vector Machine, SVM) based on pixels features.

## 1 Introduction

Human impact over Earth’s ecosystems is polymorphic. In addition to the most mediatized effects such as global warming and animal species extinction, another characteristic of the human impact on Earth’s ecosystem is the multiplication of biological invasions due to the recent globalization of human society. The interest of the scientific community for this ecological threat is rather recent too and dates from the late fifties with the publication of “The Ecology of Invasions by Animals and Plants” by Charles Elton (Elton, 1958). During the following decades, issues related to the conservation of local environments rose with the emergence of disciplines such as environmental management, biodiversity conservation and ecology restoration, for which the propagation of alien invasive plants constitutes a major problem (Zhang, 2010). This category of trees, herbs, shrubs and ferns can be described as plants capable of growing outside of their natural habitats with a stronger hability to survive, reproduce and disperse than the indigenous species, usually with identical characteristics such as high competitive capacity and increased encroachment on disturbed environments (Royimani, 2019), making them likely to fill gaps in native ecosystems and threaten native biodiversity in the worst cases (Asner, 2008). Despite a few positive impacts in terms of both economy and ecology (Shackleton, 2007; Matongera, 2016), the impacts caused by invasive species are far more destructive. Their ability to mobilize natural resources (sunlight, water, nutrient, etc) is such that they can easily supplant indigenous species and drastically reduce the local biodiversity (Gaertner, 2009). They can also alter fire frequency (Pejchar, 2009), soil properties (Hellmann, 2017), carbon, water and nitrogen cycles (Asner, 2010; Espinar, 2015; Hughes, 2005) and indirectly jeopardize human economic activity (Hestir, 2008).

Consequently, there are multiple interests in studying these invasive species, whether to simply monitor their spread or assess their impacts. Traditional mapping techniques imply ground-based measurements supported by GPS technologies which is time consuming, expensive and cannot be easily applied to very large areas (Lawrence, 2006). It also requires direct contact with the non-native species, implying the risk to contribute to its dispersal (Hestir, 2008). Remote sensing methods and technologies present many benefits likely to mitigate these issues. Remote sensing devices, especially satellites ones, are able to quickly monitor large and remote areas. Furthermore, satellite sensors can easily handle a repeated coverage of the same locations, allowing studies taking into account plant phenology and more generally variations across time. The launching of the first multispectral satellites dedicated to Earth observation dates from the 1970s with the beginning of the Landsat project. However, a broad use of these data by the plant invasion community only dates back to the mid-2000s, after the multiplication of satellite platforms made the access to remote sensing images easier and cheaper (Vaz, 2018). Several types of sensors showed promising results in alien invasive plant monitoring. Multispectral sensors are the most common, with usually less than ten and up to twenty broad spectral bands covering the visible, short-wave, middle-wave and sometimes long-wave infrared domains. They are characterized by a large swath-wide and a high revisit rate. However their poor spectral resolution makes it difficult to discriminate indigenous species from alien ones. Conversely, hyperspectral images are characterized by a high number of spectral bands, each associated to a narrow wavelength range. This high spectral resolution allows for an identification of plant species based on their biophysical properties, which makes the discrimination between different species based on the sole spectral information easier (Atkinson, 2014; Lu, 2012; Peerbhay, 2015). However, most of the hyperspectral sensors are still airborne, implying narrow swath-width and high costs, especially for the acquisition of multi-temporal datasets. In addition, the few spaceborne sensors such as Hyperion suffer from a coarse spatial resolution (Somers, 2013). Regarding active remote sensing, SAR (Synthetic-Aperture Radar) data can provide useful characteristics *per se*, such as cloud and canopy penetration (Ghulam, 2011; Chang, 2014) but are hampered by the iconic radar noise, the speckle, plus the fact that a high spatial resolution is only possible at the price of a narrow swath-wide.

This study is conducted in the context of the DIVINES project, to which contribute various teams of the Reunion University and the CIRAD on Reunion Island. The project have four main goals, which are the deployment of monitoring tools to follow the evolutions of the island’s ecosystems, the production of innovative biodiversity inventories, and finally the supply of biodiversity indicators and distribution maps highlighting their relations with climate and topography, using GIS and remote sensing tools. In particular, we are here interested in the capacity of remote sensing data and methods to improve the detection of invasive plants and their monitoring through time. The overall idea is to assess the ability of remote sensing tools to support ground-based mapping techniques by extending its scope to uncovered areas and timestamps. Two approaches have been considered, a macroscopic one and a microscopic one. The macroscopic approach aims to characterize a pixel-wise invasion rate (Fenouillas et al., 2020) using classification method applied on multispectral datasets. Meanwhile, the microscopic approach is a detection-oriented approach aiming to map the distribution first and eventually the spread of specific plant species using geolocalized in situ samples as reference data (ONF, 2020). Our work has been conducted on the basis of two main hypothesis. The first one states that when working with multispectral datasets, spectral information is not enough to discriminate properly different plant species. However, the spread of alien invasive species should create distinctive textural patterns in relation with the crown and foliage structure of a particular species easily recognizable thanks to the high spatial resolution of these data. Using textural features in macroscopic invasion rate mapping and microscopic species detection processes should then greatly improve the accuracy of the results.

In this document, the two approaches will be addressed successively, with the presentation of working and reference data, of the textural features used and the first results obtained.

## 2 Macroscopic approach

### 2.1 Working data

For this study we worked with a SPOT acquisition extracted from the Kalideos platform, a database created in 2001 by the CNES in order to promote remote sensing data. The image has been registered on July 7, 2019, over the southeast part of the Reunion island, and we considered two datasets, each associated to a specific processing level. The first one is level 1A (numerical counts including radiometric and geometric calibration) which we had to georeference ourselves and the second one is level 2A, which include orthorectification and top of canopy reflectances. We focused on a specific part of the dataset corresponding to the volcano area and its surroundings (Figure 1).

**Figure 1:**
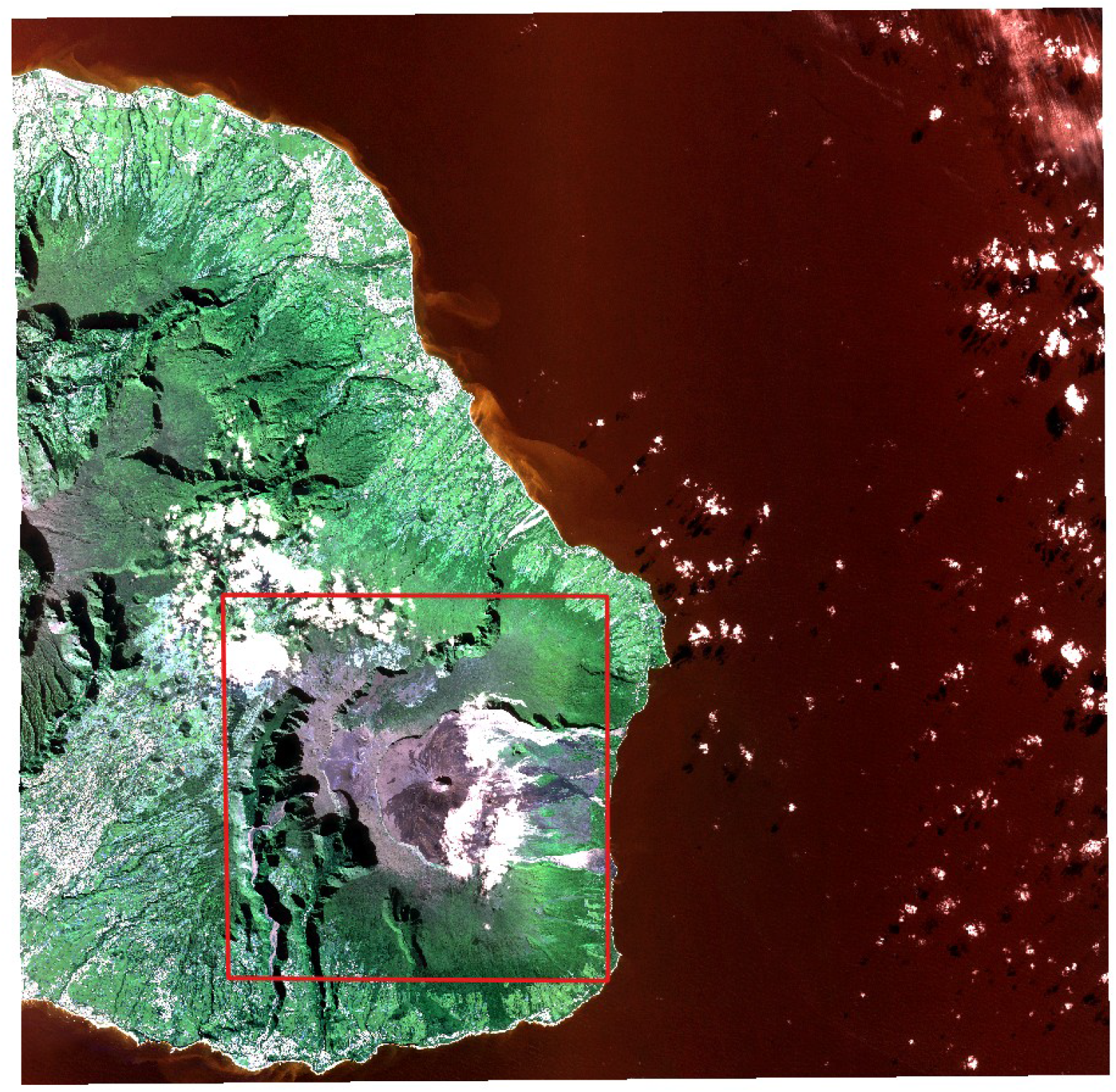
SPOT image (RGB) used in this study, with a specific focus on the area outlined in red

The ground truth dataset used in order to validate our processes is a map of the overall invasion level at island scale (Figure 2, see Fenouillas et al, 2020 for details). This map is the product of the combination of survey data gathered by several organizations (National Park of La Réunion, National Botanical Conservatory of Mascarin, ONF, DEAL) which had to be harmonized in order to constitute a unique classification map with four invasion level classes at 250 m horizontal resolution: very low/intact, low, moderate and high.

**Figure 2:**
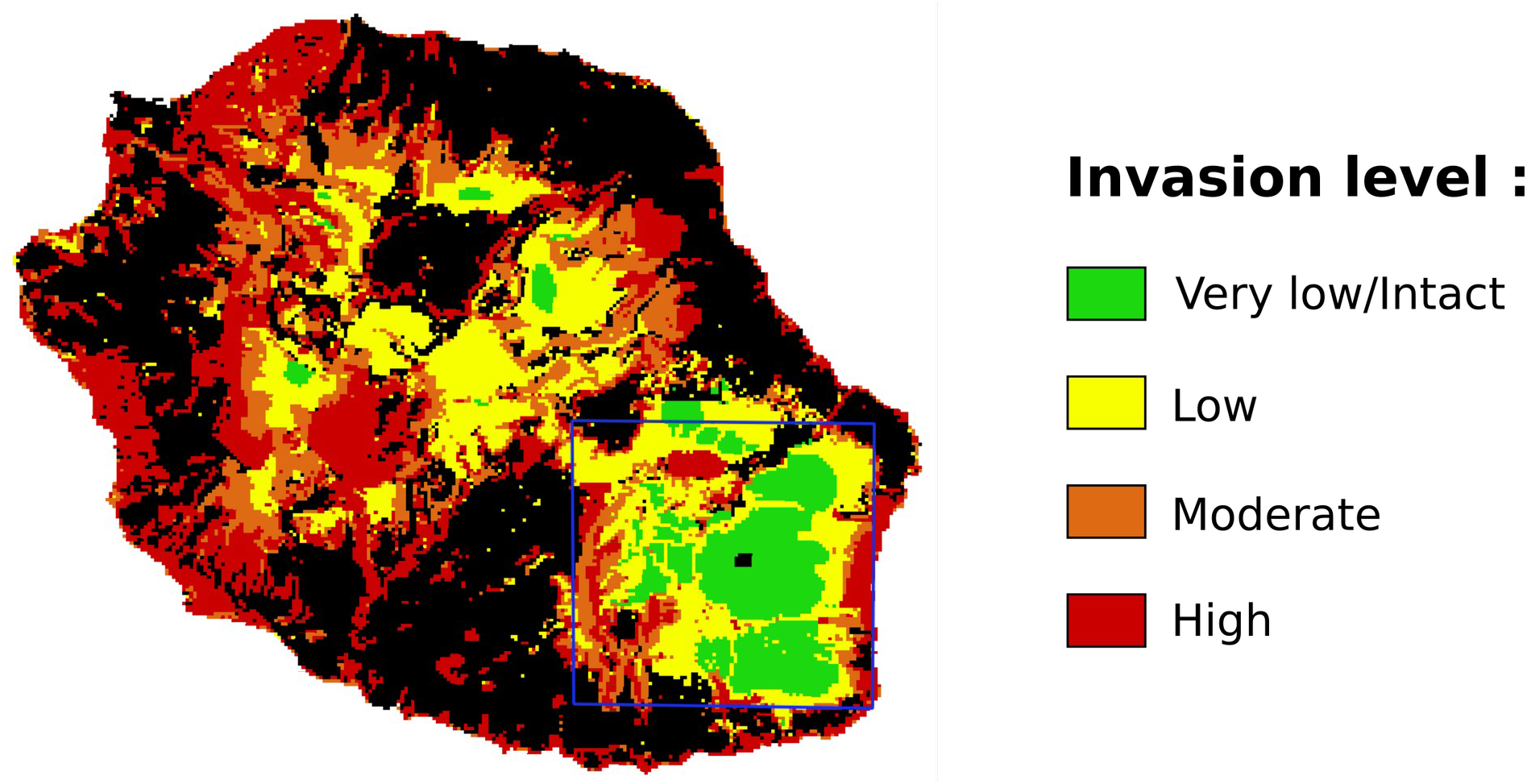
Invasion map characterized by four invasion levels (black pixels correspond to no data, Fenouillas et al. 2020)

**Figure 3:**
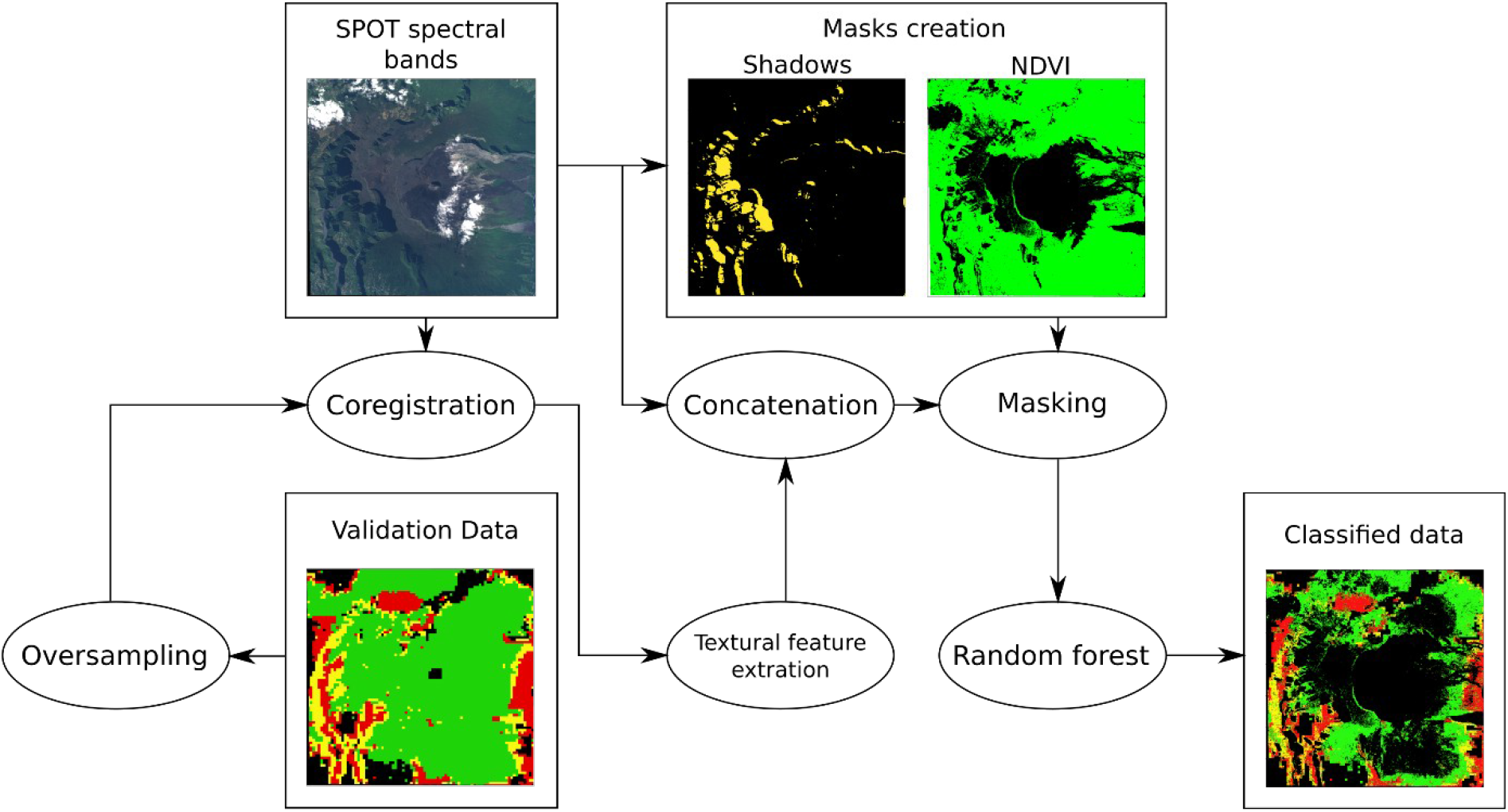
Classification workflow

Before starting the classification process, all these datasets had to be coregistered, including a geolocalization step for level 1A SPOT dataset and an oversampling step for the validation dataset so it can match the spatial resolution of each working dataset.

### 2.2 Classification method

The objective of this study is to assess the ability of the textural information to compensate for the poor spectral information provided by multispectral data in the context of indigenous/invasive plant species discrimination. Our classification framework is detailed in Figure 1. After the coregistration step, a shadow mask is built using the *c1c2c3* color space in order to ignore shadow pixels. This color space is known to present good shadow invariant properties (Salvador, 2004), and especially the *c3* component:

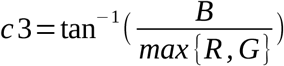

where *R, G* and *B* are respectively the red, green and blue channels of the SPOT image. The shadow mask is then obtained by thresholding this new component with a threshold value set to 0.95.

A vegetation mask is also built in order to only consider the vegetated pixels. This mask is computed by thresholding the normalized difference vegetation index (NDVI), computed as follows:

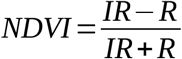

where *IR* and *R* are respectively the infrared and red components of the SPOT image. Here we consider vegetated pixels as those associated to a NDVI value superior to 0.2, which corresponds to various vegetation density, from very low canopy cover to complete canopy cover. Pixels that are not excluded by these two masks are kept as working pixels in our study. Clouds are associated to very low NDVI values and will therefore be excluded by this masking step as well as shadow and non-vegetation pixels.

In parallel of the masking step, textural features are computed from the four spectral bands of the SPOT image. In this study we considered two different features: Haralick coefficients based on the definition of a co-occurrence matrix, and morphological profiles. A cooccurrence matrix (Haralick, 1973) is a data structure presenting the proportion of pixels passing from one value to another for a given spatial shift *D* (*x, y*), and a given neighborhood (Figure 4).

**Figure 4:**
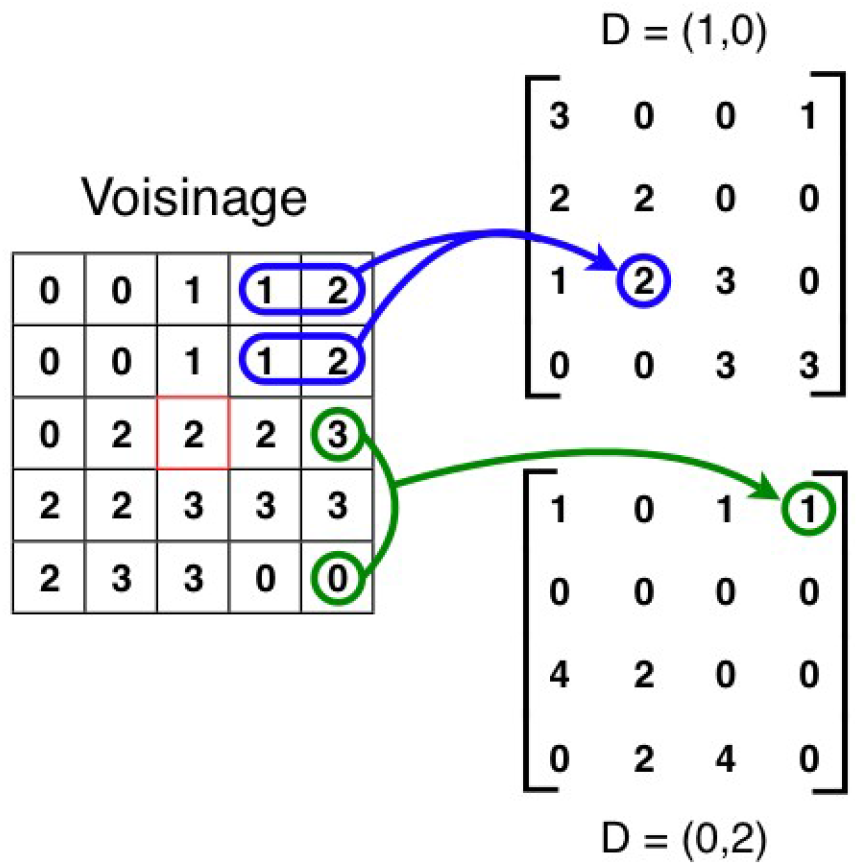
Exemple of the creation of two co-occurrence matrices, each associated to a different shift.

**Figure 5:**
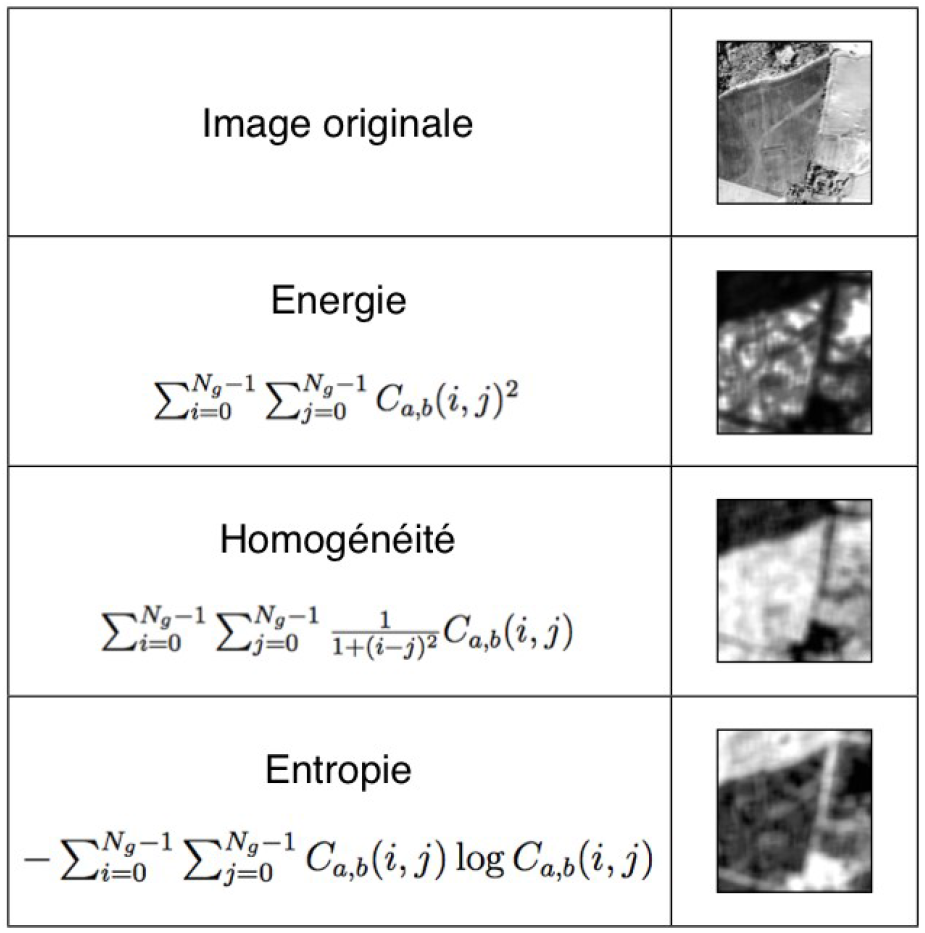
Exemples of Haralick coefficents applied on co-occurrence matrices

The size of the matrix is equal to the number of gray-levels in the image, so the dynamic of the image has to be reduced in order to avoid the creation of gigantic matrices. That’s why the data are previously undersampled to a 16 classes classification map using an improved version of the K-means algorithm (Linde, 1980). Because co-occurrence matrices are typically large and sparse, various metrics are derived from the matrix to get a more useful set of features called Haralick coefficients. These coefficients are related to specific textural characteristics of the image (homogeneity, contrast, etc) or to the complexity and nature of the gray-level transitions which occur in the image (Figure 1).

In this study, we focused on four specific coefficients: contrast, homogeneity, energy and dissimilarity. They are computed for each pixel of a spectral band using a sliding window, for four different shifts which are eventually averaged: *D*(0,1), *D*(0,−1), *D*(−1,0) and *D* (1,0). Finally, five window sizes are considered: 5, 10, 15, 20 and 25.

Image analysis based on morphological profiles was initially proposed by Pesaresi (2001) and is based on mathematical morphology operators (Soille, 2003). Morphological profiles are intrinsically linked to the concept of granulometry, which aims to analyze the statistical distribution of structure sizes in an image by applying a set of morphological operations associated to structuring elements (i.e. small window of particular shape characterizing the neighborhoods on which the operator is applied) of increasing size, in order to build features highlighting bigger and bigger spatial structures as the neighborhood expands. In this study we used the same morphological operators as Fauvel (2008), namely the opening and closing by reconstruction. These are geodesic operators which do not modify the structures unlike classical opening/closing operators but only remove those which are smaller than the structuring element. In a non-binary context, opening profiles will highlight dark structures of increasing size while closing profile will highlight bright structures of increasing size (Figure 6). Several sizes of disk-shaped structuring elements were tested to build the extended morphological profiles, from 4 to 68 pixels.

**Figure 6:**
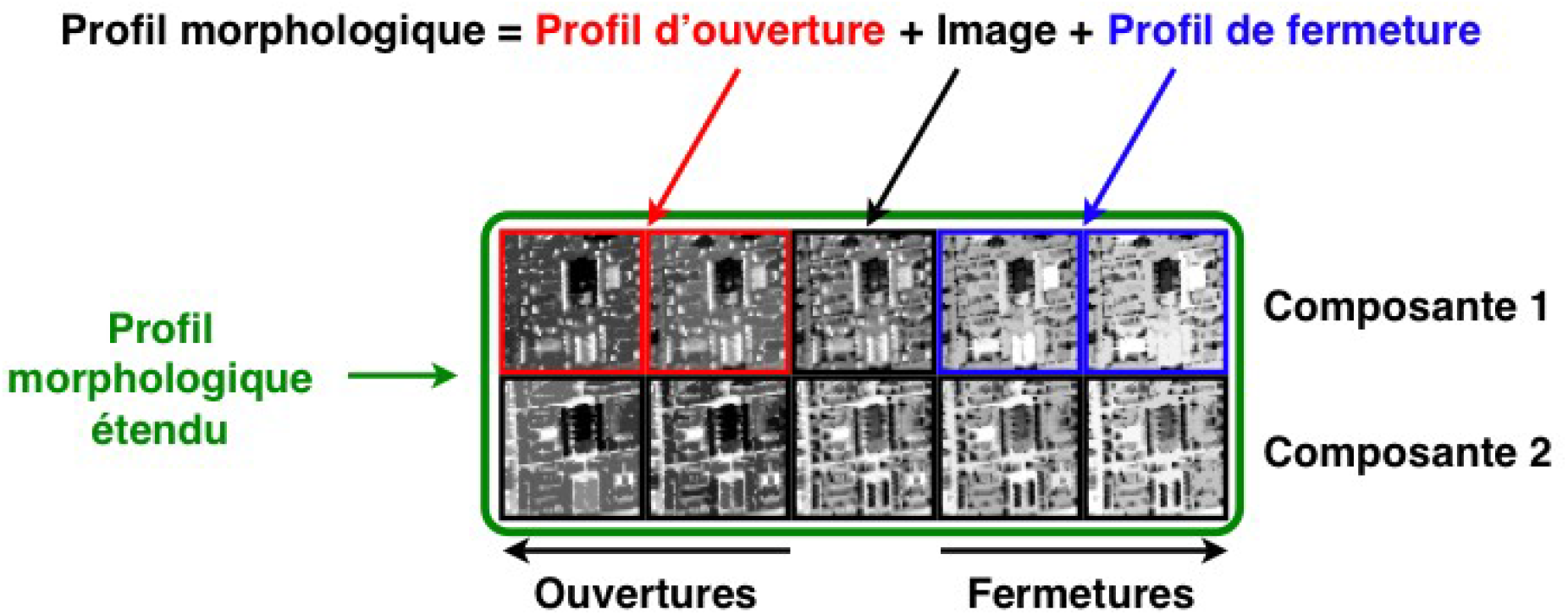
Structure of an extended morphological profile

At the end of the process, the spectral and/or textural features we want to use in order to build the classification map are concatenated, masked and set as input of the classifier, a Python implementation of the Random Forest algorithm provided by the Sklearn library. It is a supervised algorithm, which means that it has to be trained over sufficiently exhaustive and representative data in order to produce relevant outputs. The final set of valid pixels is divided in two subsets: the training set (10% of the valid pixels) and the validation set (90% of the valid pixels) with no overlap between both of them. The first one is used to produce a model which is validated on the validation set.

### 2.3 Results and discussion

The whole testing area mentioned in part 2 is large. In order to limit computing time for the following tests, we focused first on a smaller area (Figure 7) and verified in the end that we are able to get approximately the same results on the large dataset with the most efficient set of parameters.

**Figure 7:**
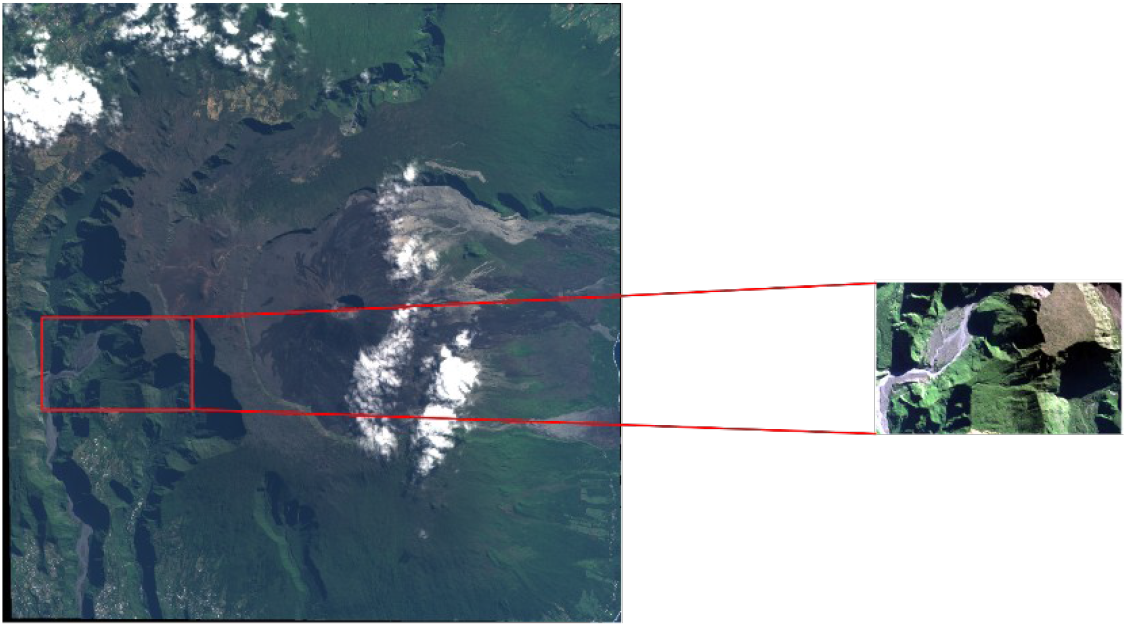
Working area for the analysis of textural features parameters

The first series of tests were dedicated to spectral features only. Table 1 shows that multispectral information only is globally inadequate to evaluate the level of invasion. Whether with one spectral feature, the whole four or principal components (new features built through principal component analysis as linear combinations of the original features maximizing the variance of the dataset), the overall accuracy never exceed 60%. However we can already remark that the extreme levels of invasion (very low and high) are systematically better classified than the intermediate ones (low and moderate).

**Table 1:** Classification accuracy obtained with spectral features only. R = red, G = green, B = blue, IR = infrared and PC = principal component

| Features | Classification accuracy (%) |  |  |  |  |
| --- | --- | --- | --- | --- | --- |
|  | Very low | Low | Moderate | High | Total |
| R | 46 | 41 | 34 | 34 | 39 |
| R + B | 60 | 46 | 33 | 54 | 49 |
| R + B + IR | 61 | 48 | 36 | 53 | 50 |
| R + G + B + IR | 65 | 54 | 41 | 68 | 57 |
| 4 PC | 69 | 54 | 44 | 70 | 59 |

With textural features concatenated to the spectral ones, the results are far better. Table 2 shows that depending on the number of textural features given as input to the classifier, the results can be up to 30 % more accurate than with spectral features only. It also shows that morphological profiles can quickly improve the accuracy of the classification even with a few small structuring elements but haralick coefficient computed with large windows lead in the end to better overall accuracies. The last line of the table finally shows that the fusion of the two intermediate classes is another way to improve the classification results, which means that these two classes are maybe too similar to each other.

**Table 2:**
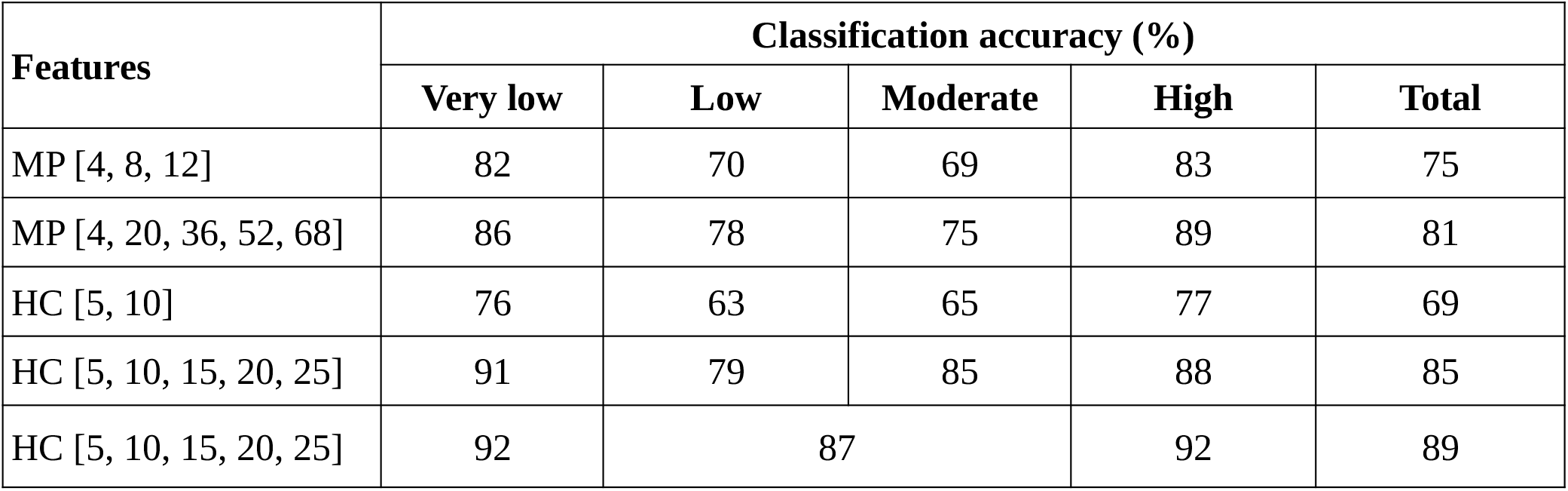
Classification accuracies obtained with spectral and textural features. MP = morphological profiles (the numbers indicate the structuring elements used to built the profiles) and HC = Haralick coefficients (the numbers indicate the window size used to build the co-occurrence matrices)

All these results were obtained on level 1A SPOT data. Applied to level 2A SPOT data, results are systematically more accurate, with improvements ranging from two to seven percent compared to the less preprocessed data.

## 3 Microscopic approach

### 3.1 Working data

For this approach, we worked on the same SPOT 6 dataset than in the previous section, but we focused on three different sub-areas (Figure 8). A reference dataset composed of several polygons corresponding to camphor trees (*Cinamomum camphora*, Lauraceae) and Cryptomerias (*Cryptomeria japonica*, Taxodiaceae), two species easily recognizable at canopy level, was provided by nationala forestry services (ONF, 2020). It is worth noting that only the camphor trees are considered as invasive. Cryptomerias are not recognized as invasive, but will be used as a second easily recognizable reference species.

**Figure 8:**
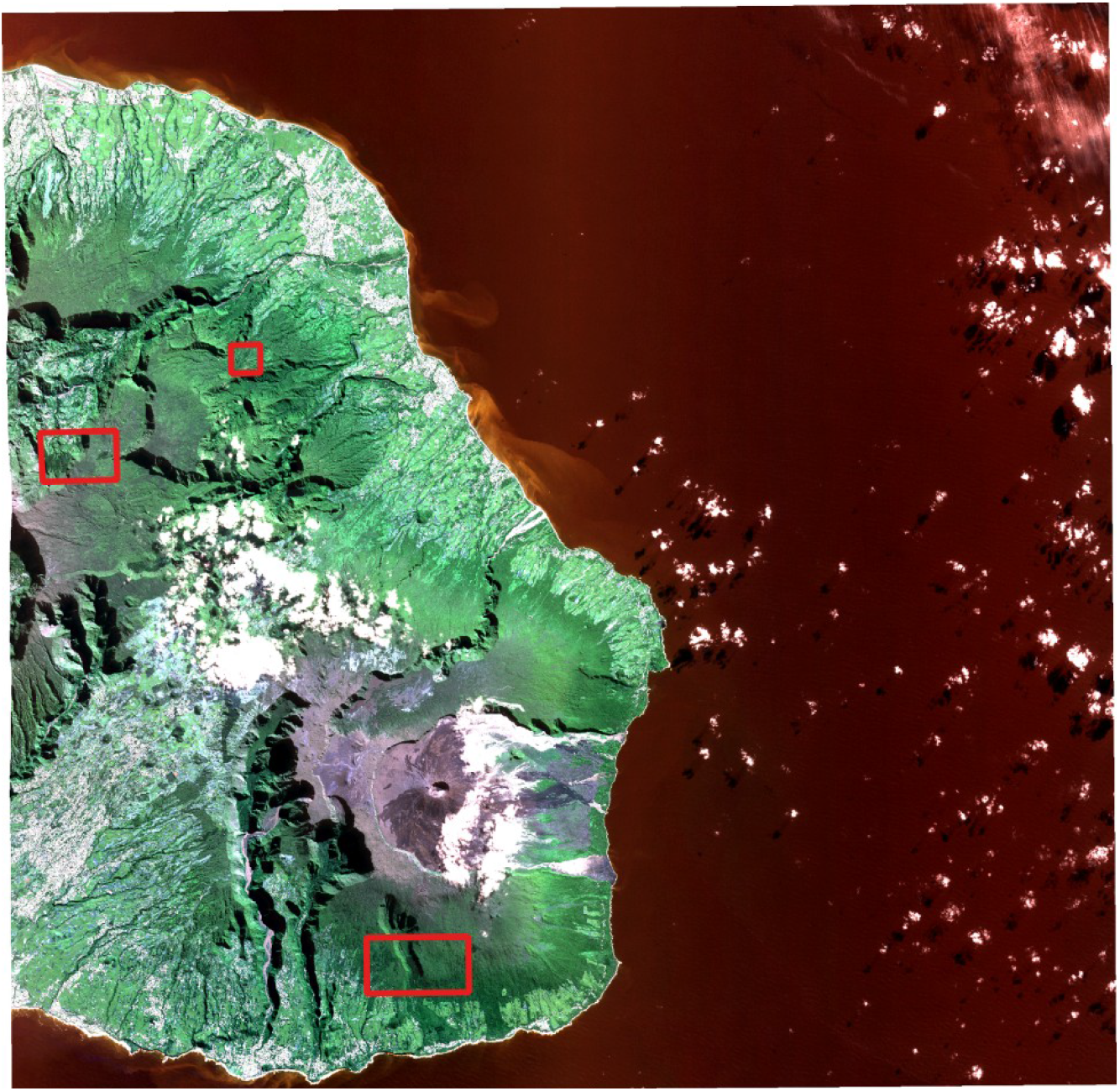
SPOT dataset with the three new working sub-areas outlined in red

Each sub-area includes several polygons corresponding to planted sites of camphor trees and/or cryptomerias without shadow or cloud obstacles (Figure 9). The number of samples for each species and each sub-area are listed in Tableau 3.

**Figure 9:**
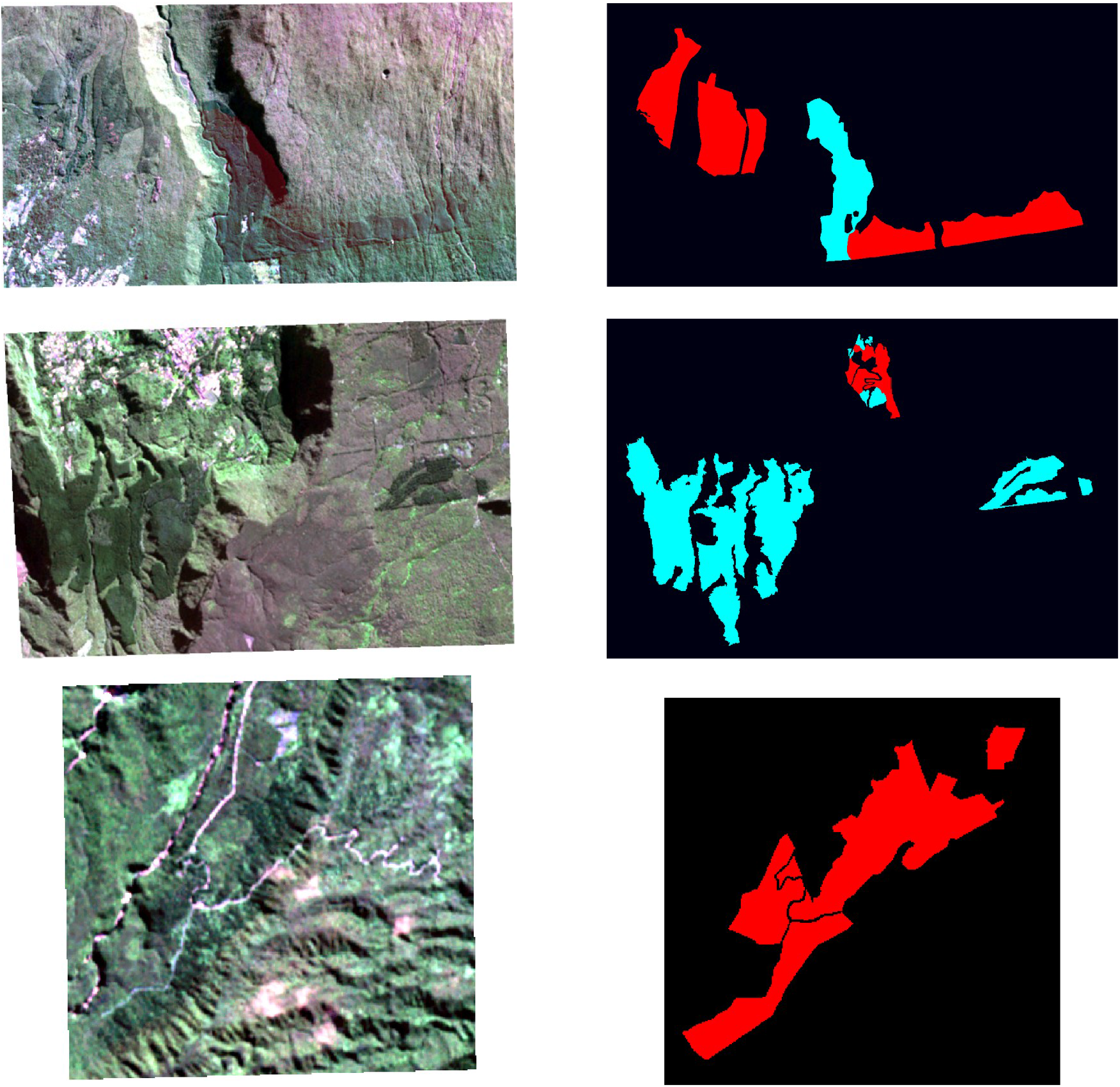
The three areas of interest (SPOT data on the left, ground truth on the right), from area 1 on the top and area 3 on the bottom. Red polygons correspond to camphor habitats and cyan polygons to cryptomeria habitats (ONF, 2020).

**Tableau 3:** Number of samples for each species and each sub-area

| Species | Area 1 | Area 2 | Area 3 |
| --- | --- | --- | --- |
| Camphor | 67132 | 4213 | 15926 |
| Cryptomeria | 23552 | 50410 | - |

### 3.2 Classification method and results

Several monoclass classification methods have been tested: One-class SVM, Isolation forests and Local Outlier Factor detection. These are supervised approaches for which 10% of the samples have been used for the learning step and 90% for the test step. One-class SVM produced by far the best results, so we will focus on this method only. Two quality criteria are considered:

- Precision (P) is the proportion of true positive (TP) samples over the whole number of positive samples, true and false (FP): P = TP / (TP + FP)
- Recall (R) is the proportion of true positive samples over the whole number of true samples: R = TP / (TP + TN)

The following tables show the results obtained with spectral features only (Table 4) and spectral/textural features (Tables5 and 6).

**Tableau 4:** Detection results obtained for the specific microscopic approach with spectral features only, with precision in red (P) and recall in green (R)

| Species | Area 1 |  | Area 2 |  | Area 3 |  |
| --- | --- | --- | --- | --- | --- | --- |
|  | P | R | P | R | P | R |
| Camphor | 0.27 | 0.40 | 0.53 | 0.23 | 0.42 | 0.16 |
| Cryptomeria | 0.36 | 0.42 | 0.85 | 0.41 | - | - |

**Tableau 5:**
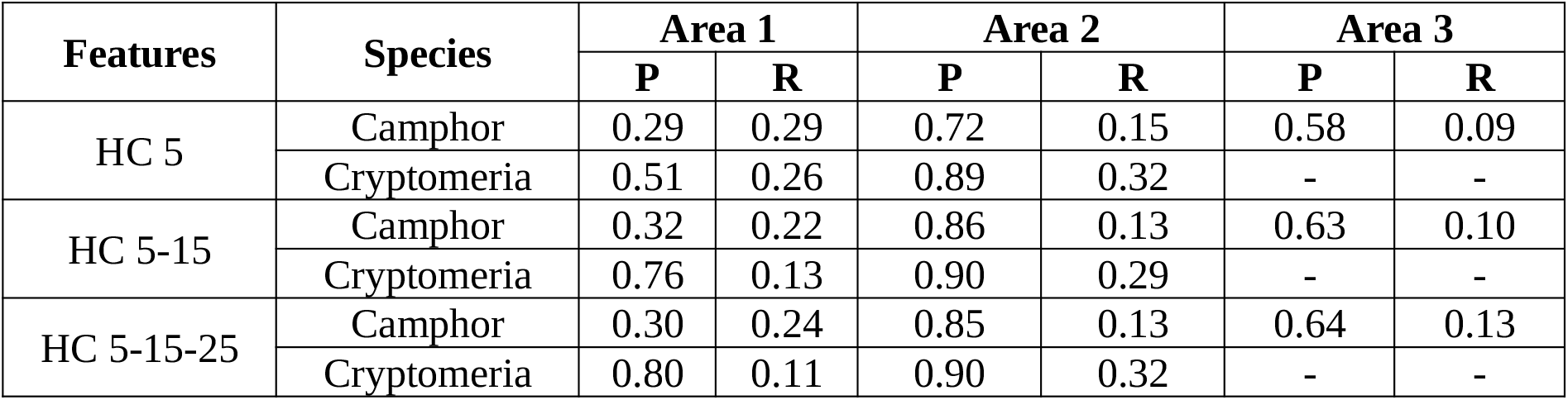
Detection results obtained with spectral / textural features (Haralick coefficients), with precision in red (P) and recall in green (R)

| Features | Species | Area 1 |  | Area 2 |  | Area 3 |  |
| --- | --- | --- | --- | --- | --- | --- | --- |
|  |  | P | R | P | R | P | R |
| HC 5 | Camphor | 0.29 | 0.29 | 0.72 | 0.15 | 0.58 | 0.09 |
|  | Cryptomeria | 0.51 | 0.26 | 0.89 | 0.32 | - | - |
| HC 5-15 | Camphor | 0.32 | 0.22 | 0.86 | 0.13 | 0.63 | 0.10 |
|  | Cryptomeria | 0.76 | 0.13 | 0.90 | 0.29 | - | - |
| HC 5-15-25 | Camphor | 0.30 | 0.24 | 0.85 | 0.13 | 0.64 | 0.13 |
|  | Cryptomeria | 0.80 | 0.11 | 0.90 | 0.32 | - | - |

**Tableau 6:** Detection results obtained with spectral/textural features (Morphological profiles), with precision in red (P) and recall in green (R).

| Features | Species | Area 1 |  | Area 2 |  | Area 3 |  |
| --- | --- | --- | --- | --- | --- | --- | --- |
|  |  | P | R | P | R | P | R |
| MP 8 | Camphor | 0.58 | 0.19 | 0.56 | 0.23 | 0.83 | 0.08 |
|  | Cryptomeria | 0.68 | 0.24 | 0.93 | 0.25 | - | - |
| MP 8-12 | Camphor | 0.70 | 0.18 | 0.53 | 0.21 | 0.86 | 0.06 |
|  | Cryptomeria | 0.66 | 0.24 | 0.95 | 0.22 | - | - |
| MP 8-12-20 | Camphor | 0.71 | 0.18 | 0.54 | 0.22 | 0.77 | 0.07 |
|  | Cryptomeria | 0.73 | 0.25 | 0.96 | 0.23 | - | - |
| MP 8-12-20-36 | Camphor | 0.71 | 0.18 | 0.44 | 0.22 | 0.77 | 0.10 |
|  | Cryptomeria | 0.73 | 0.26 | 0.96 | 0.24 | - | - |
| MP 8-12-20-36-52 | Camphor | 0.65 | 0.21 | 0.51 | 0.23 | 0.65 | 0.10 |
|  | Cryptomeria | 0.73 | 0.29 | 0.96 | 0.25 | - | - |

Spectral-only results are not very accurate, in terms of both precision and recall. The only exception concerns the cryptomeria on area 2 for which the detection precision is quite good (Table 4). Adding textural information to the detection process has various effects depending on the species, the area and the type of feature used. First, it must be noticed that the use of textural features systematically decreases recall rates by a factor of up to four, but overall increases precision, in particular for Camphor. Haralick coefficients greatly increase the precision of the detection of cryptomeria trees on area 1, but were nearly useless for the detection of camphor trees. On area 2, results are opposite while on area 3, a slight improvement can be noticed for the detection precision of camphors. When these features are effective, they are all the more effective as the scale of the coefficient is high. Morphological profiles slightly increase the precision of the detection for all three areas and, for both species (Table 6). However, this improvement does not increase with the scale of the textural features, it may even decrease. We note however that the quality of the spatial data regarding the actual distribution of the focal species is relatively poor. Better results should be obtained with better resolved and more accurate field data.

## 14 Conclusion and perspectives

In this study, two approaches for the analysis of the spread of invasive vegetal species using remote sensing data were addressed. The first one is a classification framework using both spectral and textural information to discriminate several overall qualitative levels of alien plants invasion. We showed that in this context, the presence of textural information is crucial to produce relevant classification results. The second one is a detection method focused on detecting particular plant species. It aims at detecting a single known class among an unknown number of other classes using the very same spectral and spatial features as the first approach. Unfortunately, the results of this second approach are much less promising. Even if the textural information generally increases the precision of the detection algorithm, it comes at the price of a drastic decrease of recall. It means that the algorithm is less often wrong, but also that the proportion of undetected valid samples is higher.

A first perspective to this work would be to test the generalization capacity of the Random Forest classification models created in this study over datasets acquired on different timestamps, especially for top of canopy data (level 2A). Second, we could test how much the phenology could improve the two approaches by building models based on several datasets acquired throughout the same year. A third possibility would be to investigate the potential of ITC (Individual Tree Crown) characterization methods on high-resolution data (Pléiades) and finally, to replace the One-class SVM used for the second approach by a convolutional neural network.

